# The onset time of balance control during walking is phase-independent, but the magnitude of the response is not

**DOI:** 10.1101/655928

**Authors:** Hendrik Reimann, Tyler Fettrow, David Grenet, Elizabeth D. Thompson, John J. Jeka

## Abstract

The human body is mechanically unstable during walking. Maintaining upright stability requires constant regulation of muscle force by the central nervous system to push against the ground and move the body mass in the desired way. Activation of muscles in the lower body in response to sensory or mechanical perturbations during walking is usually highly phase-dependent, because the effect any specific muscle force has on the body movement depends upon the body configuration. Yet the resulting movement patterns of the upper body after the same perturbations are largely phase-independent. This is puzzling, because any change of upper-body movement must be generated by parts of the lower body pushing against the ground. How do phase-dependent muscle activation patterns along the lower body generate phase-independent movement patterns of the upper body? We hypothesize that in response to a perceived threat to balance, the nervous system generates a functional response by pushing against the ground in any way possible with the current body configuration. This predicts that the changes in the ground reaction force patterns following a balance perturbation should be phase-independent. Here we test this hypothesis by disturbing upright balance using Galvanic vestibular stimulation at three different points in the gait cycle. We measure the resulting changes in whole-body center of mass movement and the location of the center of pressure of the ground reaction force. We find that the whole-body balance response is not phase-independent as expected: balance responses are initiated faster and are smaller following a disturbance late in the gait cycle. Somewhat paradoxically, the initial center of pressure changes are larger for perturbations late in the gait cycle. The onset of the center of pressure changes however, does not depend on the phase of the perturbation. The results partially support our hypothesis of a phase-independent functional balance response underlying the phase-dependent recruitment of different balance mechanisms at different points of the gait cycle. We conclude that the central nervous system recruits any available mechanism to push against the ground to maintain balance as fast as possible in response to a perturbation, but the different mechanisms do not have equal strength.

## Introduction

The control of balance during walking for humans is an important problem because the failure to maintain balance leads to falls and often injury. In the United States alone, costs from fall-related injuries amount to 20-30 billion USD annually (Burns, Stevens, & Lee, 2016). The upright human body is mechanically unstable, with a high center of mass and a relatively small base of support. Balancing this body is already a challenge during standing, requiring continuous regulation of muscle activity by the central nervous system (Morasso & Schieppati, 1999). As a result of intensive research over the last decades, we understand this neural control of balance during standing reasonably well (e.g. Kiemel, Zhang, & Jeka, 2011; Maurer, Mergner, & Peterka, 2006; Peterka, 2002; Winter, 1995).

Balance control during walking has been studied much less in comparison. One reason for this is that walking is highly nonlinear, with the configuration of the body changing substantially as contact with the ground is established and lost at different points during the gait cycle. This implies that at different points in the gait cycle, the central nervous system has different options available for how to generate force along the body and against the ground to affect balance. One well-studied mechanism is to shift the location of the foot placement when taking a step, which changes the pull of gravity on the body during the subsequent swing (Bruijn & Van Dieën, 2018; Hof, 2008; Reimann, Fettrow, & Jeka, 2018; Wang & Srinivasan, 2014). Another mechanism is to use ankle musculature to pull on the body during single stance (Hof & Duysens, 2018; Hof, Vermerris, & Gjaltema, 2010; Reimann, Fettrow, Thompson, & Jeka, 2018). Compared to foot placement shift, the ankle mechanism is limited in effect by the relatively small area of contact under the stance foot. Moments across the ankle joint both pull the body sideways and the foot up, so excessively large moments will result in the foot rolling over. On the other hand, the ankle mechanism has the advantage of being able to act much faster throughout single-stance and even double stance, whereas the foot placement mechanism can only be used when taking a step. For optimal benefit, there is preliminary evidence that humans flexibly coordinate these two balance mechanisms (Fettrow, Reimann, Thompson, Grenet, & Jeka, 2018), and possibly others, to adaptively respond to different challenges of balance during walking.

Responses to balance-related perturbations generally depend upon the phase of the gait cycle in which they are applied. Reflexes in the lower leg, probed by cutaneous electric stimulation, are highly modulated during walking, and even reverse direction (Yang & Stein, 1990; Zehr & Stein, 1999). Changes of foot placement in response to electric stimulation of the vestibular system are also highly phase-dependent (Bent, Inglis, & McFadyen, 2004). In contrast, responses in the upper body to these stimulations do not, or only weakly, depend on the phase of the perturbation. Logan et al. (2014) found similar results in response to visual perturbation, where the lower body response was strongly phase-dependent and the upper body response only weakly. Since the only way to affect the movement of the upper body is to generate a muscle force that acts on the lower body and the ground, these results pose the question of how the central nervous system achieves this phase-independence of the upper body.

We hypothesize that the central nervous system flexibly coordinates different balance mechanisms in the lower body in a phase-dependent way to stabilize the movement of the whole body in a phase-independent way. Here we test this hypothesis experimentally by perturbing humans walking on a treadmill with Galvanic vestibular stimulation at three different points in the gait cycle, and analyzing how the response in the whole body center of mass (CoM) and the functional force response depend upon the phase of the perturbation. We define the functional response as the displacement between the center of pressure (CoP) and the CoM. This variable is proportional to the acceleration of the CoM when assuming a single-link inverted pendulum model of the body biomechanics (Hof, Gazendam, & Sinke, 2005), which has proven to be reasonable approximation in a variety of cases (Kuo, 2007). We expect that the functional balance response to these stimuli does not depend on the phase of the perturbation. We focus our analysis to the frontal plane, which is more challenging for balance control compared to the sagittal plane (O’Connor & Kuo, 2009).

## Methods

Twenty young, healthy subjects (ten female) volunteered for this study. Subjects were between 19 and 38 years old (22.2 ± 4.57), 170.7 ± 8.4 cm tall and weighed 70.5 ± 13.5 kg. Subjects provided informed verbal and written consent to participate. Subjects with self-reported history of neurological disorder or surgical procedures involving the legs, spine or head were excluded. The experimental design was approved by the Temple University Institutional Review Board.

### Experimental Design

The study was conducted in the virtual reality setup of the Coordination of Balance and Locomotion laboratory at Temple University, which we described in detail in Reimann, Fettrow, Thompson, and Jeka (2018). Subjects walked on a split-belt treadmill in a virtual environment projected onto a curved dome that covered almost their entire field of vision (Bertec, Inc.). The treadmill was self-paced, using a nonlinear PD-controller in Labview (National instruments Inc., Austin, TX, USA) to keep the markers on the posterior superior iliac spine on the anterior-posterior mid-line of the treadmill. The same speed command was sent to each belt of the treadmill. The virtual environment consisted of a tiled marble floor with floating cubes randomly distributed in a volume 0-10 m above the floor, 2-17 m to each side from the midline, and infinitely into the distance, forming a 4 m wide corridor for the subjects to walk through (see Figure 1), implemented in Unity3d (Unity Technologies, San Francisco, CA, USA). The perspective in the virtual world was linked to the midpoint between the two markers on the subject’s temples, superposed over the forward motion defined by the treadmill speed.

**Figure 1.**
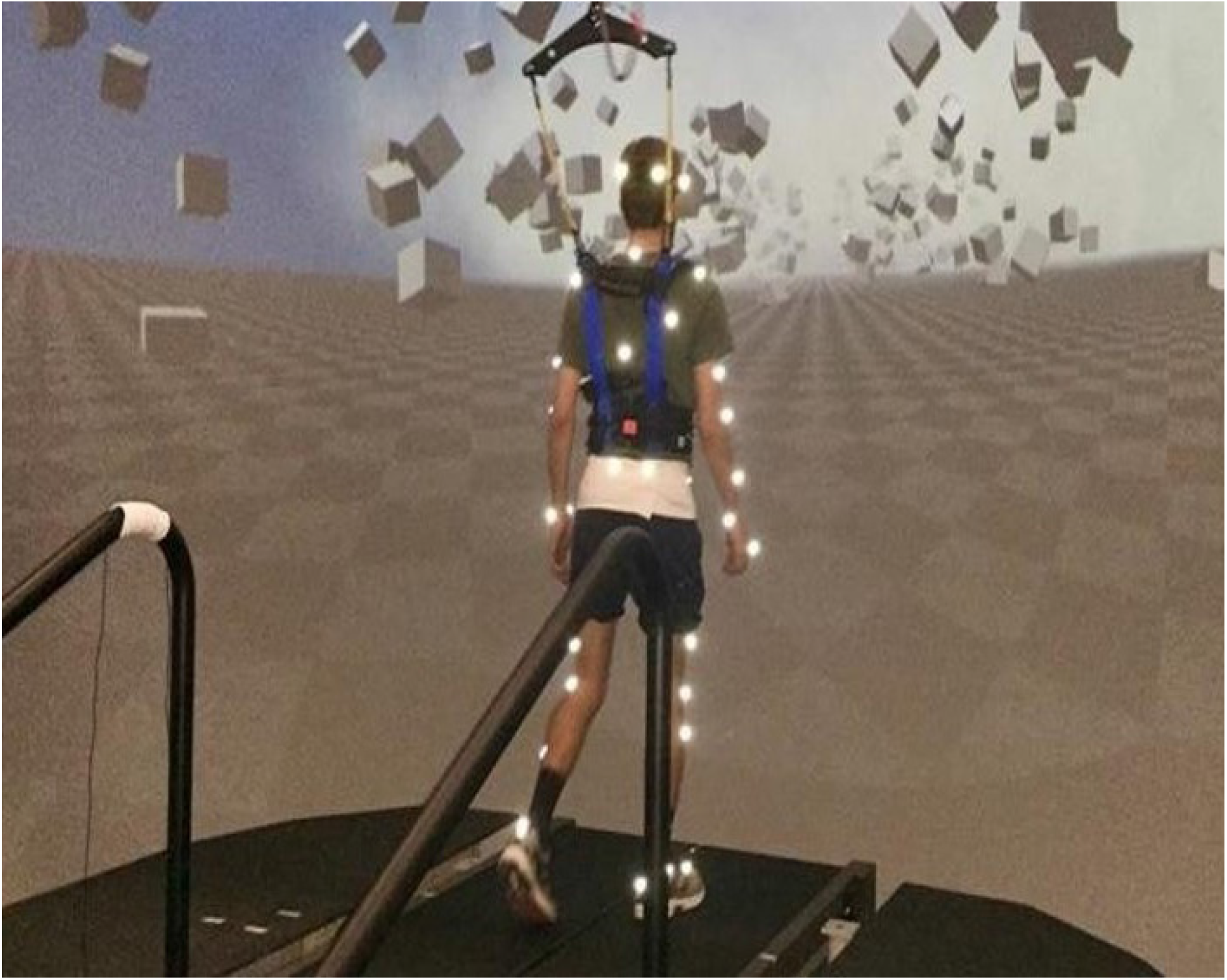
Experimental setup. Subjects walked on a self-paced treadmill immersed in a virtual environment projected onto a curved screen that covered almost their complete field of vision. The perspective in the virtual world was linked to the subject’s head position.

We used Galvanic vestibular stimulation (GVS) to induce the sensation of a fall to the side. Stimuli consisted of a 500 *μ*A current between two round electrodes (3.2 cm diameter, Axelgaard Manufacturing 103 Co., Ltd, Fallbrook, CA, USA) fixed to the mastoid processes behind the ears. The stimuli were triggered on heel-strikes of the right foot and lasted for 600 ms. Between the triggering heel-strike and the stimulus onset, we added a randomized delay of 0, 150 or 450 ms, which we will refer to as EARLY (0 ms), MID (150 ms) or LATE (450 ms) stimulus. The polarity of the stimulus was randomized to induce the sensation of falling either to the RIGHT or LEFT. A randomized wash-out period of 6-8 strides followed each stimulus. Heel-strikes were defined as downward threshold crossings of the vertical heel-marker position, where the threshold was set to 3 mm above the vertical heel-marker position of each foot during quiet standing.

Forty-five reflective markers were placed on the subject, using the Plug-in Gait marker set (Davis, Ounpuu, Tyburski, & Gage, 1991) with six additional markers on the anterior thigh, anterior tibia, and 5th metatarsal head of each foot. Marker positions were recorded at 250 Hz using an infrared motion capture system with 9 cameras (Vicon, Inc). We collected surface electromyographical data bilaterally from five muscles along the legs and hips, but the data was corrupted due to problems with the measurement device (Delsys, Inc), precluding analysis. Ground reaction forces and moments were collected at 1000Hz from both sides of the split-belt treadmill and transformed into a common coordinate frame to calculate whole-body CoP (Winter, 1990).

After explaining the experiment, obtaining consent and placing markers and EMG sensors, subjects first walked for 15 minutes on the self-paced treadmill in the virtual environment to adapt to this experimental setup. We then stopped the treadmill and exposed subjects to the GVS while standing to familiarize themselves with the sensation, then asked them to respond to this balance perturbation “normally” while walking on the treadmill. Data collection blocks consisted of two alternating phases for *metronome* and *stimulus*. During metronome phases, lasting 30s, subjects were provided an auditory metronome at 90 beats per minute and asked to use this as an “approximate guideline” for their pace, both during metronome and stimulus phases. During stimulus phases, lasting 120 s, the metronome was turned off, and subjects received GVS as described above. Data were collected only during stimulus phases. Each subject performed four blocks of walking, each block consisting of five metronome and five stimulus phases, always starting with metronome phases, for a total of 12.5 minutes per block. After each block, the treadmill was stopped and subjects were offered a break. This protocol was implemented in a custom Labview program that sent the head position and treadmill speed to the Unity computer via UDP and the stimulation currency to the stimulator.

### Data Processing

We filled small gaps of up to 100 ms length in the kinematic data using cubic splines, then low pass filtered with a 4th order Butterworth filter at a cut-off frequency of 10 Hz. From the marker data, we calculated joint angle trajectories based on a geometric model with 15 segments (pelvis, torso, head, thighs, lower legs, feet, upper arms, forearms, hands) and 38 degrees of freedom. We estimated the hip joint centers based upon pelvis landmarks (Bell, Pedersen, & Brand, 1990; Tylkowski, Simon, & Mansour, 1982) and the knee joint centers and knee flexion rotational axes from reference movements using the symmetrical axis of rotation approach (Ehrig, Taylor, Duda, & Heller, 2007). We performed inverse kinematics by minimizing the distance between the measured and the model-determined marker positions (Lu & O’Connor, 1999). This optimization was performed first for the six pelvis degrees of freedom, which formed the root of the kinematic tree, then for the six degrees of freedom at the lumbar and cervical joints, and last for each of the arms and legs separately. We estimated the body center of mass (CoM) position based on estimated segment CoM locations (Dumas, Chèze, & Verriest, 2007) and the inverse kinematics and calculated CoM velocities and accelerations using numerical derivation by time. Force plate data was low pass filtered with a 4th order Butterworth filter at a cut-off frequency of 50 Hz.

We identified heelstrike events for each foot by finding minima in the vertical positions of the heel markers with inter-peak distances > 250 ms and peak prominence > 2 cm, and pushoff events as the first peak in the vertical velocity of the 2nd metatarsal marker with a prominence > 0.35 m s^−1^ after each heelstrike. We visually inspected the result of this automatic identification and applied manual corrections in the rare cases where events were misidentified. We then partitioned the data into strides that started and ended with a right heel-strike. For each stimulus, we extracted data from the stride before and after the triggering right heel-strike, a total of two strides per trigger. The stride after the triggering heel-strike was analyzed as the stimulus data, and the stride before the trigger was used as the unperturbed reference. For each stride, we time-normalized the data for each step between consecutive heel-strikes. Strides containing missing kinematic data were excluded from further analysis. After removing these strides, an average of 43.2 ± 6.8 stimulus strides remained for each subject.

For the *whole*-*body response*, we analyze the velocity and acceleration of the CoM. Although these are not independent of each other, we chose to include both, because the acceleration is more directly related to the forces applied by the muscles, but the velocity is less susceptible to processing artifacts from numerical derivation. We represent the *functional response* by the displacement between CoP and CoM, which is proportional to the CoM acceleration when approximating the body by a single-link linear inverted pendulum (Hof et al., 2005). We characterized the *lower*-*body response* by changes of the swing foot placement relative to the stance foot at each step, and the *upper*-*body response* by the head and trunk roll angle. The roll angle was defined as the angle between the segment vector and the vertical in the frontal plane, where the head segment was the vector from the seventh cervical vertebra marker to the mid-point of the two markers on the back of the head, and the trunk segment was the vector from the mid-point of the two markers on the right and left posterior superior iliac spine to the seventh cervical vertebra marker. For each subject, we subtracted the mean of the unperturbed reference data from the stimulus data to estimate the response induced by the sensory perturbation. For visualization purposes, we also estimated the 95% confidence intervals for each trajectory across all repetitions, assuming no correlation from repeated measures within subjects. This estimate was only used to generate the shaded areas in the figures.

### Estimation of Response Onset Time and Magnitude

Reliably estimating the onset time of the response is difficult due to the high amount of natural variability in walking. Our approach was to extract a short interval of data starting at the stimulus onset for each trigger, where we can assume that there is no change initially, but the response starts at some point during this interval. We approximated the time-dependency of each variable over this interval by fitting a piece-wise linear model in R (R Core Team, 2013) with two segments and a variable break-point (Muggeo, 2008), separately for each of the twenty subjects and six combinations of phase delay and stimulus direction. To estimate the response onset time, we used the location of the break-point between the two linear segments. To estimate the response magnitude, we used the absolute slope of the second linear segment. Note that this variable is the slope of a velocity response and has units of m s^−2^, but since it is the slope of a linear model fit with specific constraints, we chose to not refer to it as acceleration.

The model fitting process was based on the following choices and assumptions. We assumed that there is no response initially, followed by a change of the outcome variable in a pre-determined direction after the onset time. This direction of change was expected to be the same as the fall stimulus direction for the CoP-CoM displacement, and the opposite for the CoM velocity and acceleration. To reflect these assumptions, we constrained the slope of the first linear segment to 0, and treated cases where the confidence interval for the break-point or slope included 0, or where the slope had the incorrect sign as missing data. The length of the analysis interval was 450 ms for the CoP-CoM displacement and 750 ms for the CoM velocity and acceleration. This value was chosen to be long enough to encompass the initial response to each stimulus, but short enough to avoid the more complex later modulations of each variable (see Figures 2, 3).

**Figure 2.**
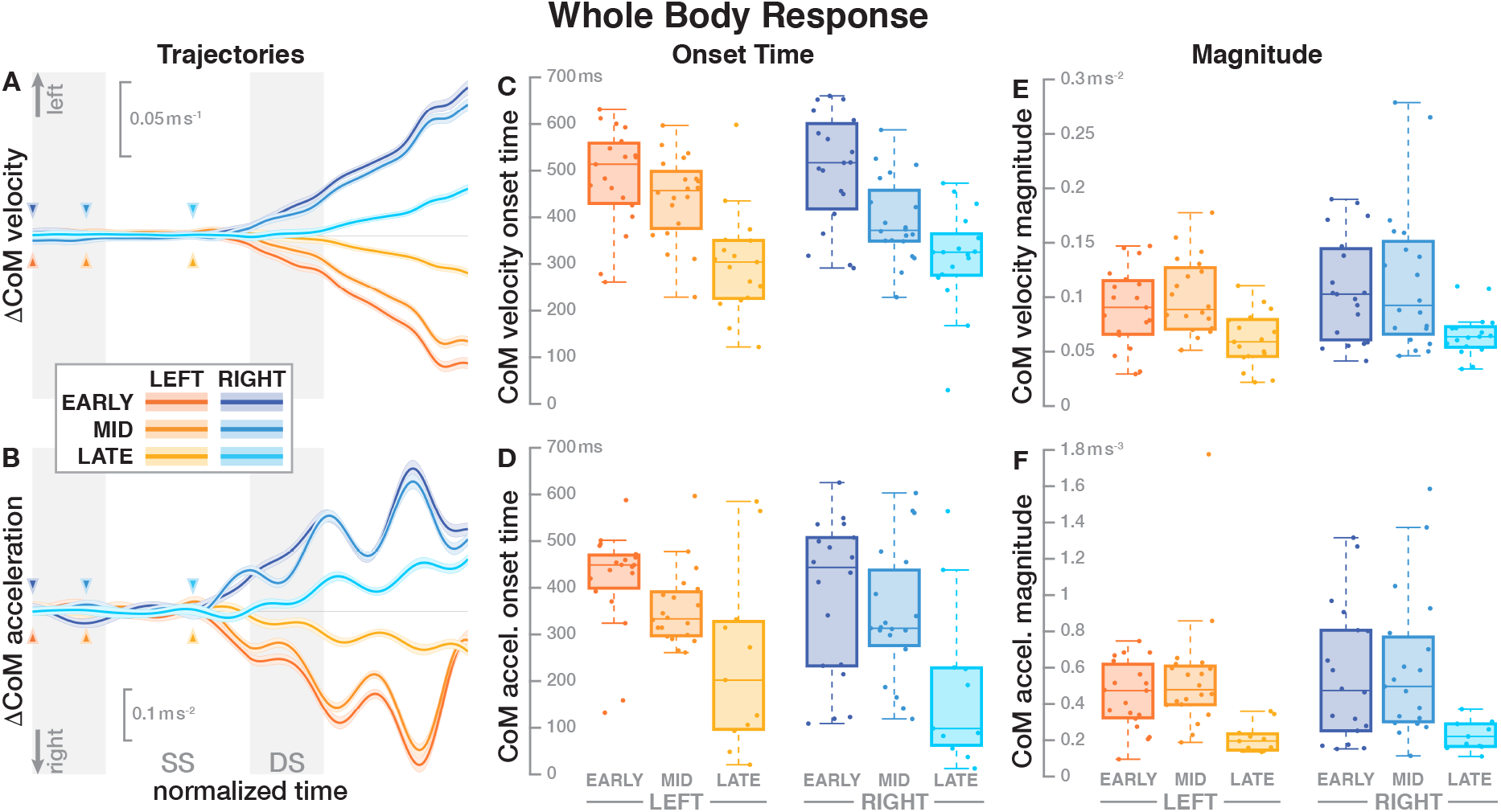
Responses in the movement pattern of the whole-body medial-lateral center of mass velocity (top row) and acceleration (bottom row). The left column shows changes in the CoM kinematics following a perturbation. Arrows mark the approximate perturbation onset. Time is normalized, showing the two steps containing and following the perturbation, with double-stance periods shaded grey and single-stance periods white. The thick lines are average responses, the color-shaded areas are 95%-confidence intervals (see Methods). The center and right column show box plots of the estimated onset time (center) and magnitude (right) of the responses. Horizontal lines are medians, the boxes cover the first to third quartiles, whiskers the upper and lower adjacents, and the dots are single data points, where each point corresponds to one subject.

**Figure 3.**
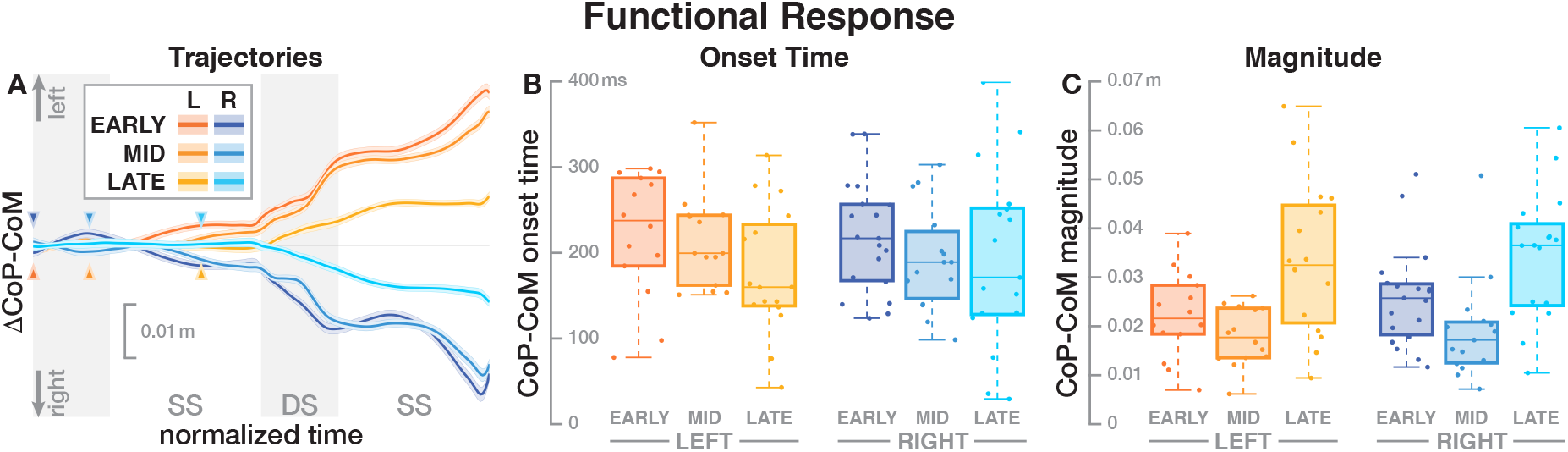
Responses in the medial-lateral center of pressure representing the hypothetical functional response to the balance perturbation. Panel A shows changes in the CoP location following a perturbation. Arrows mark the approximate perturbation onset. Time is normalized, showing the two steps containing and following the perturbation, with double-stance periods shaded grey and single-stance periods white. The thick lines are average responses, the color-shaded areas are 95%-confidence intervals (see Methods). Panels B and C show box plots of the estimated onset time (B) and magnitude (C) of the responses across subjects. Horizontal lines are medians, the boxes cover the first to third quartiles, whiskers the upper and lower adjacents, and the dots are single data points, where each point corresponds to one subject.

For the upper body variables head roll and trunk roll angle, the systematic response to the stimulus was so small relative to the natural variability from walking that the segmented model approach described above did not result in reliable estimates. For these two variables, we defined the response magnitude as the maximal excursion in the stimulus direction relative to the control average over four steps following each trigger. The lower body variable foot placement change is inherently discrete, so the notion of “onset time” is not meaningful. We defined the magnitude of the foot placement response as the change of the foot placement in the direction of the fall stimulus, i.e. the cathode.

### Statistical Analysis

For each outcome variable, we performed a linear mixed effects analysis using R (R Core Team, 2013) and *lme4* (Bates, Mächler, Bolker, & Walker, 2014), with fixed effects *phase delay* (EARLY, MID, LATE) and *direction* (LEFT, RIGHT), and random factor *subject*. We tested for significance of the fixed effects using an ANOVA with Satterthwaite’s approximation method (Fai & Cornelius, 1996), implemented in the R-package *lmerTest* (Kuznetsova, Brockhoff, & Christensen, 2017). We addressed multiple comparisons (ANOVAs for ten different variables) by Bonferroni correction. Where the ANOVA reported a significant effect, we performed post-hoc pairwise t-tests with Bonferroni-adjusted *p*-values. In all tests, we used *α* = 0.05 as significance threshold.

We initially analyzed the effect of gender as a factor in all models. Gender had no significant effect on any outcome variable (*p* > 0.05). Comparing the model with gender to a model without gender as a factor, both the Akaike information criterion and the Bayesian information criterion were consistently lower for the model without gender. Based on these results, we used the more parsimonious models without gender as a factor for further analysis.

## Results

Subjects were able to successfully complete the experimental task of walking under intermittent Galvanic stimulation. There was no instance of a subject falling or utilizing the safety harness. Subjects responded to the vestibular stimulation by swaying in the direction against the fall stimulus as expected (Reimann et al., 2017).

### Whole Body Response

Figure 2 shows the average CoM velocity and acceleration trajectories for the first two steps after the triggering heelstrike. Detailed results of the ANOVAs are reported in Table 1. For both CoM velocity and acceleration, *phase delay* had a significant effect on both the onset time and the magnitude of the response. The effect of *stimulus direction* was not significant for any upper body variable, nor was there any significant interaction between *phase* and *direction*. Panels C-D in Figure 2 illustrate these results. Both onset and magnitude of the response clearly depend upon *phase*, whereas the pattern for the two different stimulus directions is very similar. The post-hoc pair-wise t-tests for the onset of the response show that LATE is consistently different from EARLY and MID (*p* < 0.0017 for all comparisons), while MID and EARLY are significantly different from each other for CoM velocity (*p* = 0.0072), but not acceleration (*p* = 0.3230). For the magnitude of the response, LATE is consistently different from EARLY and MID (*p* < 0.0015 for all comparisons), but there is no significant difference between EARLY and MID (*p* = 1 for velocity, *p* = 0.9283 for acceleration).

**Table 1.**
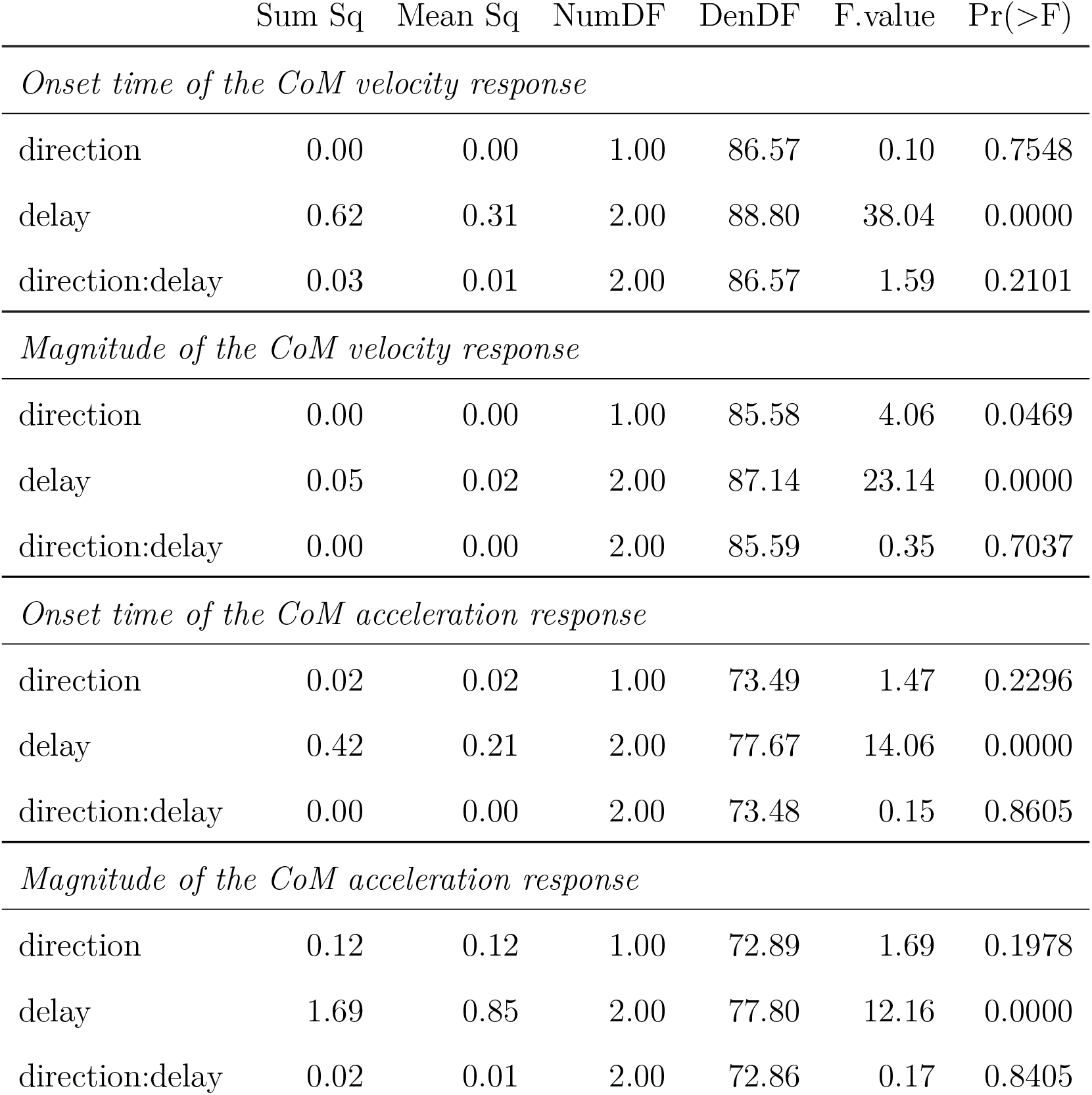
Anova results for CoM velocity and acceleration

### Functional Response

The CoP relative to the CoM shifted in the direction of the perceived fall, shown in Figure 3A. The onset time of this shift did not significantly depend upon the phase delay of the stimulus (see Table 2). Panel B in Figure 3 shows the estimated onset times. Visual inspection suggests a tendency for perturbations later in the step to have shorter onset times, but this was not significant. The magnitude of the CoP-CoM shift depended significantly upon the phase delay (see Table 2). Panel C in Figure 3 shows that the magnitude of the response was larger for LATE than for EARLY and MID for both stimulus directions, and this increase is statistically significant (post-hoc pairwise t-test, *p* < 0.0024 for both comparisons). The magnitude for MID is smaller from EARLY for both stimulus directions, but this difference is not statistically significant (*p* = 0.2135).

**Table 2.**
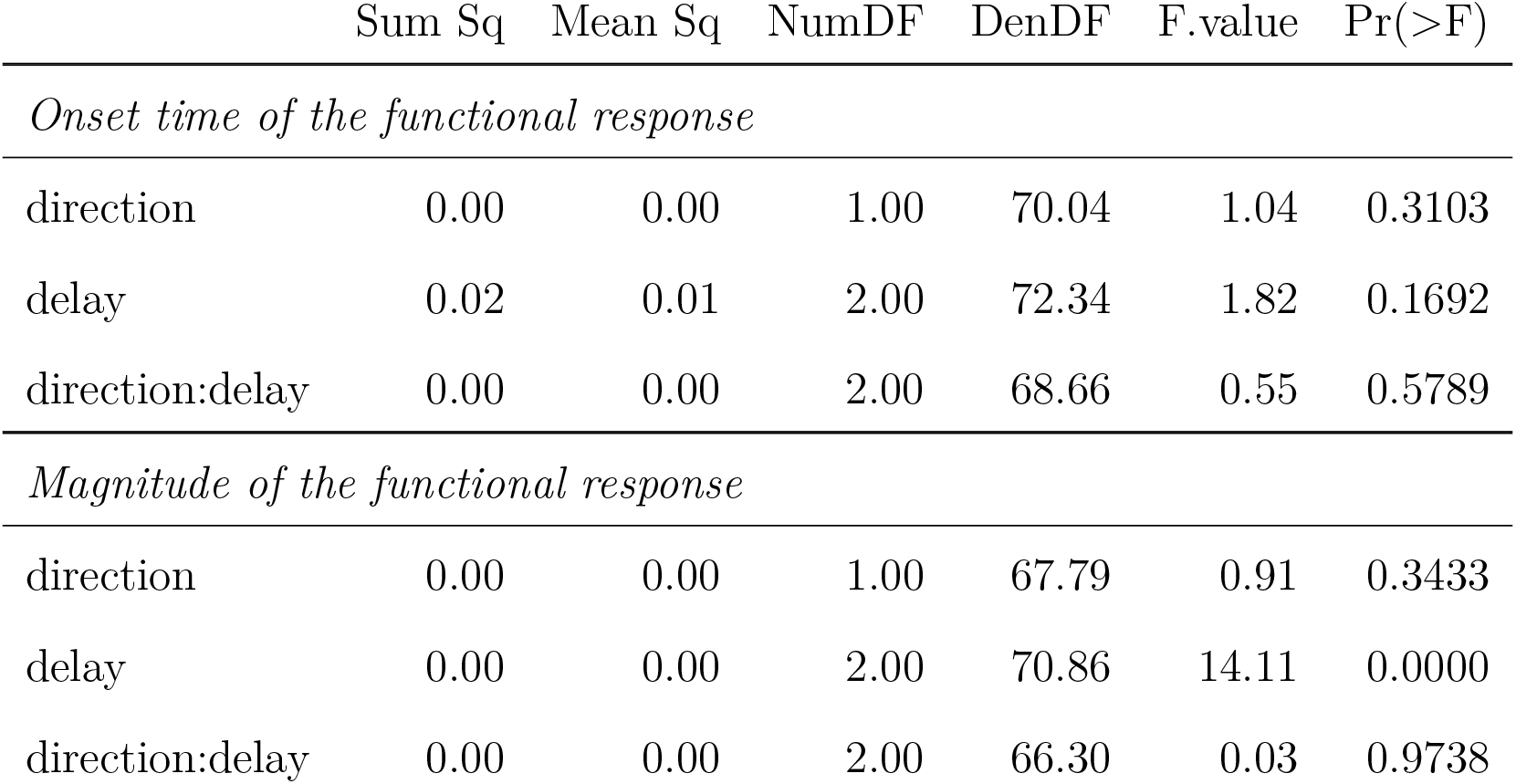
Anova results for onset time and magnitude of the function response (the CoP-CoM displacement).

### Upper Body

Both the head and the trunk segments leaned in the direction against the perceived fall in response to the stimulus. Panels A and B in Figure 4 show the average trajectories of the head and trunk roll angle for the first four steps after the triggering heelstrike, and Panels C and D show box-plots for the magnitude of the response. For the head angle, the magnitude of the roll response does not depend upon the direction, nor the phase delay of the stimulus, in a statistically significant way (see Table 3). For the trunk angle, both phase delay and stimulus direction have a statistically significant effect (see Table 3). The post-hoc pair-wise t-test shows that trunk roll magnitude is significantly different between all combinations of delays (*p* < 0.0037 for all comparisons).

**Table 3.** Anova results for magnitude of the response in head and trunk roll angle

|  | Sum Sq | Mean Sq | NumDF | DenDF | F.value | Pr(>F) |
| --- | --- | --- | --- | --- | --- | --- |
| *Magnitude of the head roll response* |  |  |  |  |  |  |
| direction | 1.41 | 1.41 | 1.00 | 5150.82 | 0.39 | 0.5330 |
| delay | 31.92 | 15.96 | 2.00 | 5149.00 | 4.39 | 0.0125 |
| direction:delay | 3.66 | 1.83 | 2.00 | 5149.06 | 0.50 | 0.6047 |
| *Magnitude of the trunk roll response* |  |  |  |  |  |  |
| direction | 224.35 | 224.35 | 1.00 | 5149.76 | 138.60 | 0.0000 |
| delay | 106.06 | 53.03 | 2.00 | 5148.61 | 32.76 | 0.0000 |
| direction:delay | 338.75 | 169.37 | 2.00 | 5148.64 | 104.63 | 0.0000 |

**Figure 4.**
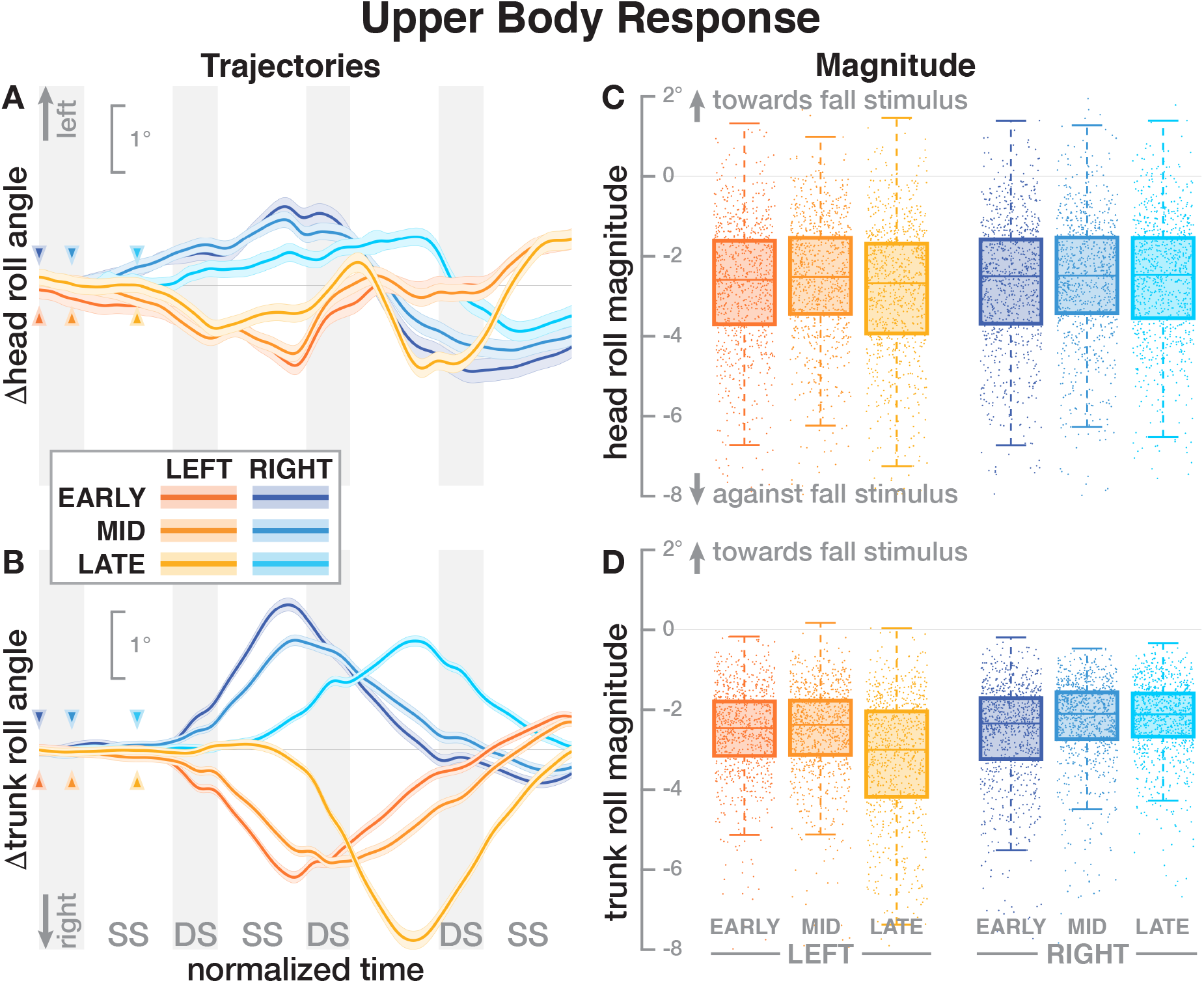
Responses in the roll angles of the head (top row) and trunk (bottom row) segments. The left column shows changes in the roll angles following a perturbation. Arrows mark the approximate perturbation onset. Time is normalized, showing the four steps containing and following the perturbation, with double-stance periods shaded grey and single-stance periods white (note that the time shown here is twice as long as in Figures 2, 3 and 5). The thick lines are average responses, the color-shaded areas are 95%-confidence intervals (see Methods). The right column shows box plots of the estimated magnitude of the responses. Horizontal lines are medians, the boxes cover the first to third quartiles, whiskers the upper and lower adjacents, and the dots are single data points, where each point corresponds to one step.

### Foot Placement

The foot placement at the fist post-stimulus step was shifted in the direction of the perceived fall, but only for the EARLY and MID phase delays (see Figure 5). At the second post-stimulus step, the foot placement was strongly shifted in the opposite direction for EARLY and MID. In the LATE condition, neither the first nor the second foot placement seems to have shifted. The left side of Figure 5 show the trajectories of both heel markers for the first two steps following the trigger, and the right side shows box plots for the magnitude of the foot placement change. Table 4 reports the results of the ANOVAs for magnitude of the foot placement change. Phase delay has a significant effect on both the first and the second step. Stimulus direction is not significant on the first step, but is significant on the second step. There is no interaction between delay and direction. The post-hoc pairwise t-tests show that foot placement is significantly different between all combinations of delays on both post-stimulus steps (*p* < 0.0001 for all comparisons).

**Figure 5.**
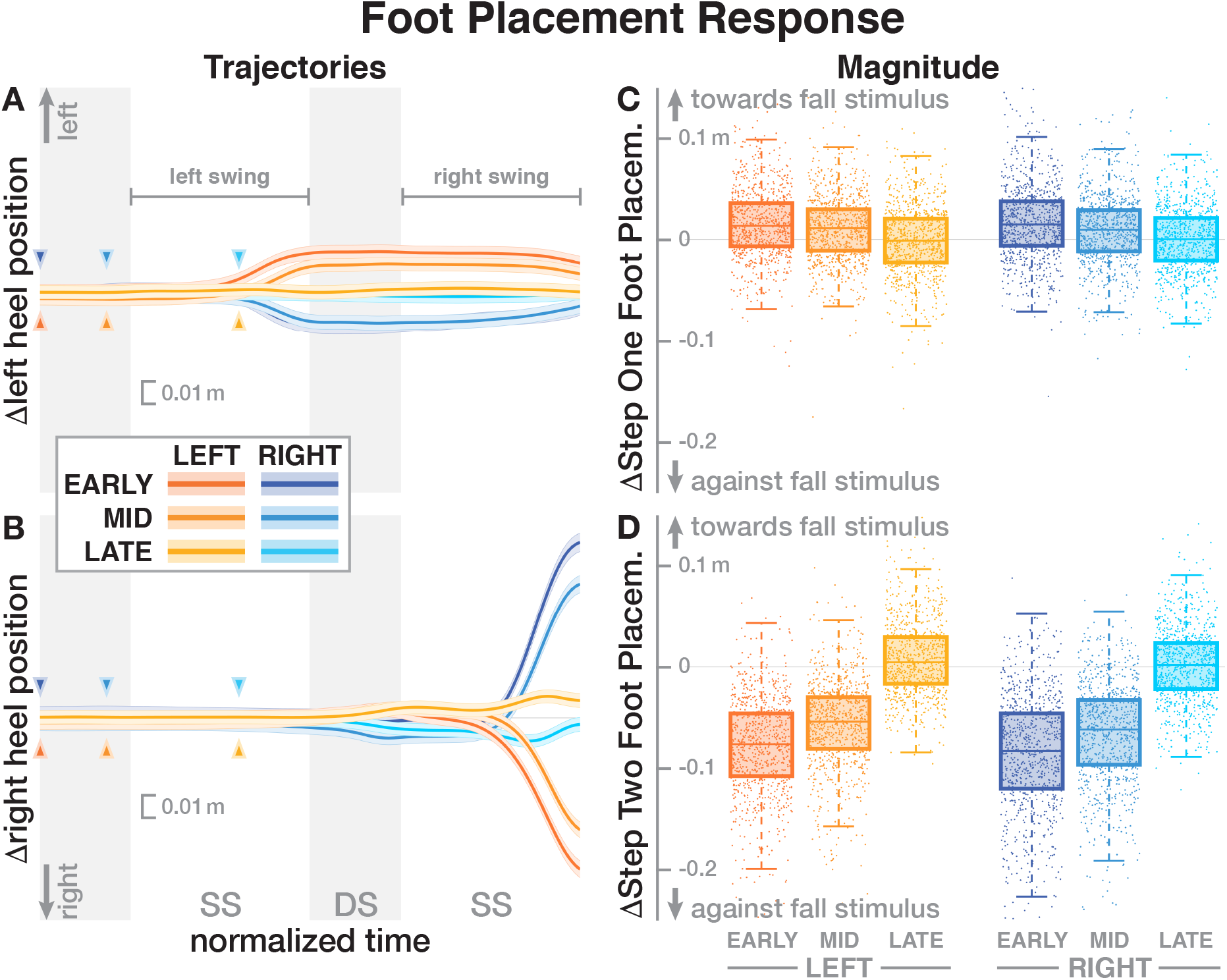
Responses of the medial-lateral foot kinematics. The left column shows changes in the average medial-lateral heel position for both legs. Arrows mark the approximate perturbation onset. Time is normalized, showing the two steps containing and following the perturbation, with double-stance periods shaded grey and single-stance periods white. The thick lines are average responses, the color-shaded areas are 95%-confidence intervals (see Methods). The right column shows box plots of the estimated and magnitude of the foot placement response, i.e. the medial-lateral position of the leading foot heel relative to the trailing foot heel at each heel-strike. Step one (C) corresponds to the time in the middle of the trajectories on the left. Step two (D) corresponds to the time at the end of the trajectories on the left. Horizontal lines are medians, the boxes cover the first to third quartiles, whiskers the upper and lower adjacents, and the dots are single data points, each point corresponds to one step.

**Table 4.** Anova results for magnitude of the foot placement response the two post-stimulus steps

|  | Sum Sq | Mean Sq | NumDF | DenDF | F.value | Pr(>F) |
| --- | --- | --- | --- | --- | --- | --- |
| *Magnitude of the foot placement response in step ONE* |  |  |  |  |  |  |
| direction | 0.00 | 0.00 | 1.00 | 5177.21 | 0.20 | 0.6551 |
| delay | 0.24 | 0.12 | 2.00 | 5168.41 | 100.97 | 0.0000 |
| direction:delay | 0.00 | 0.00 | 2.00 | 5168.39 | 1.00 | 0.3672 |
| *Magnitude of the foot placement response in step TWO* |  |  |  |  |  |  |
| direction | 0.08 | 0.08 | 1.00 | 5162.46 | 46.39 | 0.0000 |
| delay | 7.20 | 3.60 | 2.00 | 5161.56 | 2013.28 | 0.0000 |
| direction:delay | 0.00 | 0.00 | 2.00 | 5161.56 | 1.33 | 0.2633 |

## Discussion

We used Galvanic vestibular stimulation to perturb the sense of balance in walking humans at three different phases in the gait cycle. We analyzed the whole-body response to this sensory perturbation in the form of changes of the center of mass (CoM) movement, and the displacement of the center of pressure (CoP) relative to the CoM, which is one aspect of the ground reaction forces generating the whole-body movement changes. Based on previous studies, we expected that the whole-body response would be phase-independent, and hypothesized that the functional response of the CoP-CoM displacement would also be phase-independent.

Our results broadly support and extend the finding that vestibular information is used to maintain balance during walking and that the balance response to vestibular stimuli depends on the phase of the gait cycle (Bent et al., 2004; Dakin, Inglis, Chua, & Blouin, 2013; Iles, Baderin, Tanner, & Simon, 2007). Bent et al. (2004) reported that upper-body kinematic responses to GVS were independent of the phase of the gait cycle but that the foot-placement response was phase dependent. Dakin et al. (2013) found that there is vestibular influence on motor output across many muscles throughout the gait cycle, but that the contribution of any individual muscle to the overall response is highly phase-dependent.

Subjects responded to the fall stimulus by shifting their CoP in the direction of the perceived fall, i.e. the cathode. The onset time of this functional response did not depend on the phase of the perturbation (see Figure 3 and Table 2). This indicates that although different balance mechanisms are recruited at different points in the gait cycle, the neural controller tends to respond as fast as possible with any available mechanism. For the EARLY and MID perturbations, the foot placement following the perturbation was shifted in the direction of the perceived fall, but the first CoP shift occurred during single stance, indicating that the controller recruits the immediately available mechanism first and does not wait until other, potentially stronger mechanisms become available.

Contrary to our expectations and to results by Bent et al. (2004), the onset time of the whole-body response did depend on the phase of the perturbation. Responses to the LATE perturbations were consistently faster than to the EARLY and MID perturbations, with no significant difference between the latter two conditions (see Figure 2 and Table 1).

The magnitude of both the initial CoP shift in the direction of the perceived fall and the whole-body sway in the opposite direction depended upon the phase of the perturbation. The whole-body response was consistently smaller for the LATE perturbation compared to EARLY and MID (see Figure 2 and Tables 1). The functional response, in contrast, was consistently larger for the LATE perturbations (see Figure 3). This is not consistent with Bent et al. (2004), who found that the size of the upper-body response did not depend on the timing of the stimulus. This may be because we used a smaller GVS stimulus than (0.5 mA vs 1–1.5 mA, Bent et al., 2004) and also different stimulus timing, with our LATE stimulus falling in between their mid-stance and toe-off stimuli.

On the whole, there were few significant differences between EARLY and MID perturbations, but both of these were consistently significantly different from the LATE perturbations. The CoM responses were faster and smaller, whereas the CoP shift responses were larger, but with unchanged onset time. This difference in response pattern is peculiar, because as noted earlier, all whole-body movement has to be generated by pushing against the ground. So how can a smaller CoP shift generate a larger CoM movement change?

One possible explanation is that these differences in the whole-body movement are generated by combinations of forces that do not affect the location of the CoP. One example of such forces is a hip-roll mechanism, where the upper body is rotated around the hip, generating a shear force that pushes the CoM sideways without shifting the CoP (Horak & Nashner, 1986; Reimann, Fettrow, & Jeka, 2018). While this hip mechanism is usually discussed in standing or the single-stance period during walking, a biomechanical equivalent exists during double stance. This would involve differences in the trunk roll angle response early after the stimulus. Figure 4 shows that there are indeed differences, but they are neither large nor conclusive. Explaining such details satisfactorily generally requires a detailed model of the whole sensorimotor control loop, including neural dynamics, muscle physiology, biomechanics and interaction with the environment. While such models exist for standing (e.g. Peterka, 2000; van der Kooij, Jacobs, Koopman, & Grootenboer, 1999), there is no comparable model to explain balance control during walking. The available models of walking are not sufficiently detailed to explain the results presented here (Geyer & Herr, 2010; Song & Geyer, 2015; Taga, 1998).

We hypothesized that the CNS generates a functional, phase-independent motor response to balance perturbations by recruiting different balance mechanisms in a phase-dependent way. Base on this hypothesis, we predicted that the CoP shift relative to the CoM in response to a sensory fall stimulus would not depend on the phase-shift in the gait cycle in which that stimulus occurred. This prediction was only partially supported by the data, in that the onset of the response was phase-independent, but the magnitude was not. These results indicate that the CNS recruits available balance mechanisms as fast as possible to respond to a sensed threat to upright stability, but the strength of the response is initially limited during single stance and only realized after a period when other mechanisms also contribute.

Such a limitation of balance responses during single stance might be the underlying reason that drives some populations with balance deficits to adapt their gait patterns in characteristic ways. For example, people with Parkinsson’s Disease tend to have shorter single-stance times than age-matched controls for similar walking speed (Morris, Iansek, Matyas, & Summers, 1994). This might be because they have problems using the ankle roll mechanism for balance control and reduce the duration of the single stance period as a coping strategy.

The details of which balance mechanisms are recruited at which point in the gait cycle are still not well understood. One striking observation from our data is that for the LATE perturbation, the onset of the average CoP shift appears to align with the transition to double stance (Figure 3). But since there is no change in foot placement (Figure 5), this CoP shift cannot be generated by the well-understood foot placement mechanism. One possibility is that the transition to double stance allows recruitment of the push-off mechanism to shift weight between the two stance legs by modulating the push-off force of the trailing limb around that transition (Kim & Collins, 2015; Reimann, Fettrow, Thompson, & Jeka, 2018). Understanding this phenomenon requires a detailed kinematic analysis that we will perform in future work.

